# Seave: a comprehensive web platform for storing and interrogating human genomic variation

**DOI:** 10.1101/258061

**Authors:** Velimir Gayevskiy, Tony Roscioli, Marcel E Dinger, Mark J Cowley

## Abstract

Capability for genome sequencing and variant calling has increased dramatically, enabling large scale genomic interrogation of human disease. However, discovery is hindered by the current limitations in genomic interpretation, which remains a complicated and disjointed process. We introduce Seave, a web platform that enables variants to be easily filtered and annotated with *in silico* pathogenicity prediction scores and annotations from popular disease databases. Seave stores genomic variation of all types and sizes, and allows filtering for specific inheritance patterns, quality values, allele frequencies and gene lists. Seave is open source and deployable locally, or on a cloud computing provider, and works readily with gene panel, exome and whole genome data, scaling from single labs to multi-institution scale.

## Introduction

The falling cost of genome sequencing has enabled the sequencing of large cohorts of patients, to interrogate genomic variation and identify underlying causes of human disease. As a result, thousands of genes have been associated with disease, our understanding of genotype-phenotype correlations has significantly advanced, many bioinformatics tools have been developed and increasing complexity is being observed across the genome.

In medicine, there has been a rapid uptake of genome sequencing for gene discovery and diagnosis within the areas of rare disease, pharmacogenomics, non-invasive pre-natal screening and cancer care (Delaney et al. 2016). Genomics allows the comprehensive identification of variation, ranging from short variants affecting several bases such as single nucleotide variants (SNVs) and short insertions or deletions (Indels), to large copy number variants (CNVs), structural variants (SVs) and chromosomal aneuploidy. However, for personalised medicine to become a reality, all genomic variation will need to be identified and integrated with current biological and clinical knowledge, in formats that are accessible to all researchers and clinicians.

Interpretation of genomic variation by researchers and clinicians remains a difficult task (Amendola et al. 2016). Bioinformatics tools are designed to discover specific types of variation, ranging from point mutations to chromosome-scale copy number changes and rearrangements. To be of value, the functional impact of all variants on their surrounding genes and other functional elements in the genome must be comprehensively reported. It is also increasingly important to merge variant types together to determine overlapping events causing disease, such as a combined structural variant impacting one allele and a nonsense variant on the other in an autosomal recessive disorder. Variant annotations such as *in silico* scores, population frequencies in healthy controls and disease-focussed databases (e.g. ClinVar, COSMIC) can inform the inferred functional impact of a variant on a gene to help predict either pathogenicity or benignancy; however, these annotations and scores are scattered across many databases, making investigating a list of variants a cumbersome process. Curating candidate gene lists related to different diseases or aspects of biology remains a standard way for investigators to restrict the genomic search space, however these require frequent updates to keep up with the ever growing literature.

To address these challenges, we developed Seave, a web-based variant filtration platform that stores, queries and annotates genomic variation of all sizes. Seave was developed from the ground up for whole genome scale data and thus handles targeted sequencing data with ease. Currently supported variant types include SNVs, Indels, CNVs, SVs and regions of homozygosity (ROH). It is designed for clinicians and researchers and requires no knowledge of bioinformatics to use. Seave allows familial filtering on standard inheritance patterns using any combination of family members, and to easily edit the family structure to handle individuals with uncertain phenotype. Variants are annotated with a number of high quality databases and *in silico* prediction scores stored within Seave, with infrastructure to update these annotations over time. Gene lists can be used to restrict the genomic search space and are stored and managed within Seave; an audit trail of every change is recorded. Seave features an account system with users and groups that allows sharing of data with only those who should access it. We highlight Seave’s features and the flow of information during a typical analysis in Figure 1. We believe our novel packaging of these features in an open source platform capable of storing and querying all sizes of genomic variation represents a significant contribution to making genomic data more accessible to those who have the biological knowledge to interpret it best.

**Figure 1:**
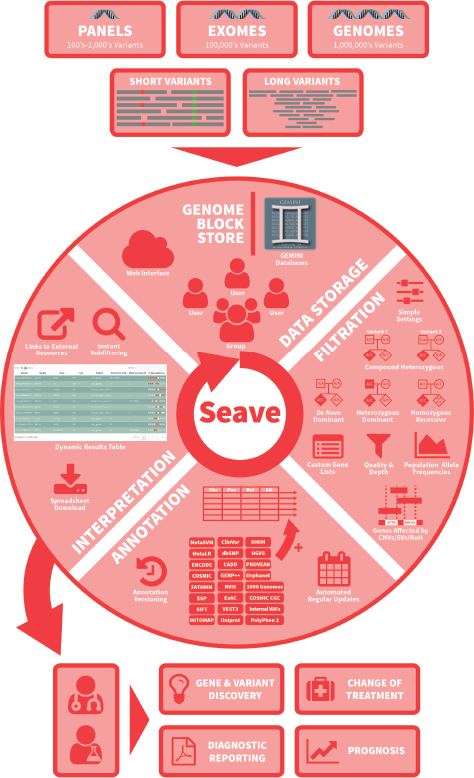
Schematic overview Seave’s main features and the flow of a typical analysis. Arrows represent the flow of information through sequencing, variant detection, data storage, filtration, annotation, interpretation and outcomes.

## Results

Seave was built at the Garvan Institute to enable gene discovery research, diagnostics and precision cancer medicine. It was predicated on a need to handle whole genome scale data and allow the flexibility to work with an ever-changing suite of bioinformatics tools for a growing list of diseases. We found existing solutions to be incapable of scale and/or flexibility and opted to build an in-house platform allowing rapid feedback from clinicians, researchers and bioinformaticians. Seave is in widespread use in Australia by multiple research groups, within clinical trials, for clinical genomics training and commercially for diagnostics.

### Rare Disease Research and Gene Discovery

Seave’s functionality enables clinicians and researchers to complement their existing biological knowledge to conduct their own variant discovery without having to learn bioinformatics. A typical scenario for rare disease diagnosis starts with genome, exome or gene panel data from an affected individual or trio and the goal of identifying the rare genomic variant(s) which segregate according to the expected inheritance pattern, within known or plausible novel disease genes, and that explain the patient’s phenotype. Pathogenic variants in the mitochondrial genome play a role in some rare genetic disorders, and Seave annotates variants with MITOMAP annotations and makes it straightforward to view the variant allele frequency as a measure of mitochondrial heteroplasmy.

Seave has been successfully used to discover novel disease genes and variants, as well as rapidly supporting the diagnosis of patients with previously reported pathogenic variants. Seave is actively used to study dozens of different rare diseases, including neuromuscular, renal, cardiovascular, mitochondrial, retinal, bone, epilepsy disorders, as well as immunodeficiencies and intellectual disability. Seave was used to elucidate the roles of *MECR* in a Mitochondrial Fatty Acid Synthesis Disorder (Heimer et al. 2016), *EPG5*-related Vici Syndrome (Balasubramaniam et al. 2017b), as well as *SLC39A8* (Riley et al. 2017) and *ECHS1* (Balasubramaniam et al. 2017a) in Leigh-like Syndromes, using a combination of exome and genome data from singletons and trios. Whole genome sequencing carried out on nine families with hereditary spastic paraplegia, and analysed in Seave, saw two patients diagnosed with novel variants in known genes and a further two diagnosed after relaxing the known disease gene list filter, leading to an expansion of the understanding of the phenotypic spectrum of mutations in *GLB1* and *PEX16* in this disease (Kumar et al. 2016). These case studies were all enabled by Seave’s inheritance filtering and rich annotations that allowed rapid discovery of known variants or gene impacts of novel variants.

### Cancer Research

The comprehensive interpretation of somatic variation represents a significant challenge. Unlike rare disease analysis, which typically focusses on finding the one or two variants responsible for a condition, there are often large numbers of damaging somatic variants of all sizes and types, and at differing variant allele frequencies (VAFs). Somatic variants of all sizes and types must be simultaneously interpreted in appropriate ways, to generate insights into more precise diagnosis, prognosis, and personalised treatment recommendations. Seave powers the analysis of somatic variation by aggregating short variants, CNVs and SVs from tumour(s) with the matched germline, if available, in one platform and connected to one another. Short variants are annotated using COSMIC and ClinVar, with numerous *in silico* predictions (Methods). A dedicated gene fusion search mode highlights potential gene fusions and the strength of supporting read evidence in terms of breakpoint quality and the number of supporting split and spanning reads.

Seave is used to study the genomic basis underlying dozens of rare and common cancer types. Seave can handle tumour-only, tumour-normal studies, and longitudinal studies investigating how tumours evolve in response to time, treatment or region. A combination of genome breadth and depth, from 150x depth WGS, to ultra-deep targeted sequencing (*>*10,000x depth panels) can be investigated. In one tumour evolution study, deep 120-150x depth WGS at multiple time points was used to investigate the genomic basis for the exceptional response to 5-azatidine therapy observed in a patient with chronic myelomonocytic leukemia (CMML) (Merlevede et al. 2016). Seave can also be used to uncover the inherited basis for cancer, and was successfully used to identify clinically relevant germline variants in 20% of patients with pituitary tumours (De Sousa et al. 2017).

Seave acts as the central hub to store and filter short variants, CNVs and SVs from the tumour and the germline when available, called from different variant callers designed for these disparate data sets. The SV fusions analysis, CNV gene list filtration, somatic annotations and regular short variant query means that Seave is well placed to handle all dimensions of a somatic research project. Collaborators are able to log in to Seave and perform queries across all variants without needing to wait for a bioinformatician, which supports rapid application of biological knowledge to the data.

### Diagnostics

Genome.One Pty Ltd is Australia’s first whole genome clinical diagnostics provider and presently uses Seave in a diagnostic capacity as part of their world’s first ISO 15189 clinically accredited laboratory for whole genome sequencing. The requirements of clinical interpretation have seen Seave thoroughly analytically validated and audited to conform to security requirements, although it does not have its own independent ISO certification. A private instance of Seave is employed with restricted firewall access to only staff members and with a subset of annotations relevant to the conditions interpreted. A team of variant interpretation specialists use Seave to filter data and, to date, 52% of over 240 cases have resulted in reportable findings being identified in genomes (Ben Lundie, personal communication).

### Precision Cancer Genomics Clinical Trials

The Lions Kids Cancer Genome Project (LKCGP) uses fast turn around whole genome sequencing of matched tumour/normal samples to improve the outcomes of kids with high-risk, or relapsed cancer where traditional treatments have been exhausted. Seave adds clinically relevant annotations (e.g. COSMIC, ClinVar), to help investigators rapidly identify the clinically relevant short variants, copy number alterations, and fusion genes for each patient. Seave has been the central hub to provide fast treatment recommendations, changes in diagnosis, and the identification of inherited cancer risk variants.

The Molecular Screening and Therapeutics (MoST) program is an innovative clinical trials framework using a targeted genomics screen to test targeted anti-cancer agents in patients with rare or advanced cancers. Using sequencing panels of 100-400 cancer genes, MoST interrogates tumour-only, or matched tumour-normal short variants and copy number variants using Seave to provide patients with a turnaround time of approximately six weeks from biopsy to results. Seave supports the analysis of about 30 cases per fortnight within this program.

### Education & Training

Seave has been used for training purposes as part of the Kinghorn Centre for Clinical Genomics’ commitment to improving genomics education. Three clinical genomics data analysis workshops with an average attendance of 60 practicing clinical geneticists, researchers, laboratory scientists and other health professionals have been run, both within Australia and internationally. Seave was extensively used to provide hands-on experience with interpreting genomic data. Particularly, inheritance filtering of variation from trios of patients provided a familiar starting point for clinicians to understand the power of genomics and how they can use it in practice. Seave was well-received and trainees found it quick and easy to use, even as cases grew progressively more complicated.

## Discussion

Precision medicine promises improved patient outcomes and better understanding of disease biology and relies on accurate identification and prioritisation of genetic variants from individual patients. Comprehensive genome interpretation will be critical to achieving this, since any sized genomic variants from single-base to entire chromosome or translocation, can be the cause of a genetic disorder or cancer. However, most variant interpretation tools only focus on SNVs and Indels, don’t scale well from targeted- to whole genome sequencing (WGS) data and commercial solutions are cost prohibitive.

Here we present Seave, a comprehensive web based variant storage, annotation and filtration platform, built to address one of the major bottlenecks in precision medicine, variant interpretation. From the millions of genetic variants identified in a patient, Seave makes it simple to store, annotate, query and filter down to the few that are relevant to the patient being studied. Seave aids in the diagnosis of patients with rare inherited genetic disorders or the identification of cancer driver mutations, as well as supporting gene discovery research and genomics education.

It does this by storing all types of variants, agnostic of their size or the method used to identify them, for small targeted-sequencing experiments, through to large WGS-sized cohorts. Variant filtering and prioritisation is enabled by a combination of frequency, quality and gene filters, high quality, up-to-date germline, somatic and mitochondrial annotations, extensive *in silico* predictions and support for family inheritance filtering.

Dedicated small variant, CNV and gene fusion queries are presented via an intuitive, secure webpage, facilitating its use by a broad range of users with no knowledge of bioinformatics required. This allows researchers and clinicians alike to leverage their knowledge, supported by Seave, to rapidly identify the variants of interest.

Variant annotations are constantly updated, making static variant annotations rapidly out of date. Seave allows administrators to update the annotation databases at a desired interval, where all annotation changes are both versioned, and reported in the result files, allowing for auditing in diagnostic settings.

Seave is designed to fit at the end of a best practice genome analysis pipeline, and as such has an API for automated data import. By leveraging the GEMINI (Paila et al. 2013) framework for storing small variants, Seave supports most variant callers, including GATK HaplotypeCaller (McKenna et al. 2010), FreeBayes (Garrison and Marth 2012), Strelka (Saunders et al. 2012) and VarDict (Lai et al. 2016). Popular CNV and SV formats are also supported, including VCF and tabular formats from Sequenza (Favero et al. 2014), CNVkit (Talevich et al. 2016), LUMPY (Layer et al. 2014), CNVnator (Abyzov et al. 2011), Manta (Chen et al. 2016) and ClinSV (Minoche et al, manuscript in preparation).

Seave runs on a very modest server and is only limited by storage space, which can easily be expanded due to the use of Amazon’s Elastic Block Storage (EBS) for storing small variants and Amazon Relational Database Service (RDS) for annotations and large variants.

The case studies presented demonstrate Seave’s broad utility for rare disease and cancer genomic applications. As CNV and SV analysis becomes more routine in genomic analysis pipelines, and WGS matures, platforms that can handle all types of genomic variants, and scale to WGS-sized datasets will be vital for the continued development of genomic medicine, and driving the next wave of disease gene discovery.

## Methods

### Analysis pipeline and importing data

Seave is a data exploration, annotation, filtration and interpretation tool and, as such, belongs at the end of a bioinformatics analysis pipeline. The primary input into Seave is short variants (SNVs and Indels) in the form of GEMINI (Paila et al. 2013) databases, which can be produced from joint-called VCF files that have been annotated with Variant Effect Predictor (McLaren et al. 2016) or SnpEff (Cingolani et al. 2012). Seave does not support importing VCFs directly as annotating WGS-sized datasets and creating GEMINI databases consumes a large amount of system resources. Any VCF that meets the VCF specification 4.1 can be used to create a database, but benefits from containing GT, GQ, AD and DP tags in the FORMAT block to make full use of Seave’s features. Seave only supports the build 37 (hg19) version of the reference genome as this is all GEMINI accommodates. Seave is tested to support VCFs from GATK HaplotypeCaller (McKenna et al. 2010), FreeBayes (Garrison and Marth 2012), VarDict (Lai et al. 2016), and Strelka (Saunders et al. 2012). Upon import into Seave, a database report is automatically generated and includes a breakdown of variant counts by categories of impact and rarity.

Seave can also import large variants such as copy number variants and structural variants from a variety of tools: CNVnator (Abyzov et al. 2011), LUMPY (Layer et al. 2014), Sequenza (Favero et al. 2014), ROHmer (Puttick et al, manuscript in preparation), ClinSV (Minoche et al, manuscript in preparation), Manta (Chen et al. 2016) and CNVkit (Talevich et al. 2016); these are imported using the unmodified outputs from each tool. We have found that even on a modest single core server, Seave scales well with databases containing variants with up to 123 whole genomes (22.3 million variants) and up to 252 exomes (3.1 million variants) for cohort-level analysis.

### Querying short variants

#### Affected status specification

GEMINI databases imported into Seave contain a fixed number of samples and short variants (Supplementary Figure 1). Seave allows classifying the samples into families, by their affected status and gender, either through a web form or pedigree file (.ped). This information is then used to enable variant filtering based on inheritance pattern in a variant-focussed manner (Supplementary Figure 2). For example, given a family with a rare disease where any number of individuals have been sequenced, a variant-focussed approach will attempt to find variants that are shared within the affected individuals in the family but not present in the unaffected individuals with the expected severity. Individuals with uncertain affected status can be easily added or removed from the family. This approach foregoes strict specification of pedigrees and is fluid to the reality of the difficulty in obtaining samples and high sequencing costs.

**Figure 2:**
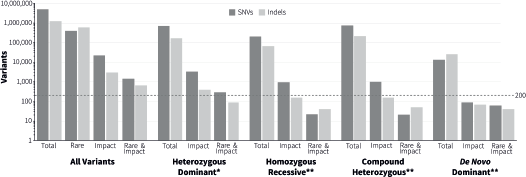
Variant counts from whole genome sequencing in the NA12878 trio restricted by combinations of rarity, gene impact (damaging) and inheritance patterns. Counts derived using the best practices GATK pipeline on raw data from the Illumina Platinum Genomes project (Eberle et al. 2017), mapped to the b37+decoy reference genome, decomposed and normalised with vt (Tan et al. 2015) and queried with GEMINI (Paila et al. 2013). Rarity is defined as a maximum allele frequency of 1% in 1000 Genomes, ExAC and ESP. Impact is defined as medium or high impact, as defined by the Ensembl impact variant annotation. *NA12878 and NA12891 were marked as affected for the purposes of this analysis. **NA12878 was marked as affected for the purposes of this analysis.

#### Inheritance patterns and families

We have implemented the standard mendelian inheritance patterns of heterozygous dominant, homozygous recessive, *de novo* dominant and compound heterozygous. Each inheritance pattern filter utilises the affected status of samples within a family to identify putative causative variants. Other inheritance patterns such as X-linked recessive can be constructed as needed, for example, by using the homozygous recessive pattern and restricting variants to chromosome X.

While a database may contain a cohort of individuals, grouping these into one or more families allows restricting sample-specific information returned to just the family selected. Combining variants from multiple families into one database allows information about variant prevalence in the cohort to be returned even when variants in just one family have been requested.

#### Filtering

After optionally selecting a family and inheritance pattern, the primary filtration happens on the query page (Supplementary Figure 3). The first filtration options are for the genomic search space of the query, this includes whether any number of genomic coordinates, saved gene lists or custom genes should be exclusively searched or excluded from the query. Exclusion of genomic search space minimises the chance of incidental findings or allows excluding known unrelated regions. Next is the impact that the variant has on a gene, categorised as low/medium/high by GEMINI (largely following the classification system of Ensembl (McLaren et al. 2016)), as well as restriction by scaled CADD score (Kircher et al. 2014). Next is the minimum acceptable prevalence of the variant in control populations of the ExAC (Lek et al. 2016), 1000 Genomes (1000 Genomes Project Consortium et al. 2015) and EVS (Tennessen et al. 2012) projects. Next are technical restrictions for the minimum sequencing depth in all samples, minimum variant quality and exclusion of variants if they have been marked as failing quality filters in the VCF. Finally, the type and maximum number of variants returned can be restricted. Using the information specified, Seave creates SQL queries used to run GEMINI on the database. Queries typically take seconds to execute but this can stretch up to a minute for cohorts of genomes with inheritance patterns specified.

#### Variant counts

A typical family trio sequenced by whole genome sequencing contains approximately 6 million short variants, however, this number is rapidly reduced when just a few of the filtration criteria described are applied. Figure 2 shows variant counts for the NA12878 trio split by SNVs and Indels across multiple filtration criteria. The total number of variants is reduced 84% by discarding variants with an allele frequency of more than 1% in population control databases or by 99.6% by discarding variants with a low impact (e.g. noncoding or silent). If both of these restrictions are applied together, 99.97% variants are discarded, leaving only 2,122. This is still too many for manual inspection, but applying inheritance patterns on top of rarity and impact results in 63 variants for the homozygous recessive inheritance pattern, 72 for the compound heterozygous pattern, 101 for the *de novo* pattern and 371 for the heterozygous dominant pattern. These counts are further reduced to single or low double digits if an intellectual disability gene list from Orphanet (Orphanet 2017), containing 1293 genes, is applied. These reductions in variant counts show that a relatively small set of criteria can be used to filter variant lists to a point where manual interpretation becomes feasible, without excluding potentially causative variants through excessive filtering.

#### Results and annotations

The results of a query are displayed on the results page in a dynamic table that includes all variants and annotations (Supplementary Figure 4). Default columns provide a summary of each variant and its impact on the genome, while individual annotation sources can be shown alongside by clicking buttons below the table (Supplementary Figure 5). The default Impact Summary column combines the major annotations into a single set of coloured icons denoting whether each annotation classifies the variant as pathogenic (red), benign (white) or does not contain any information (grey). Where an annotation source has an external web resource for the variant or gene, a link is provided that takes the user to the record, for example, clicking the genomic coordinates of the variant in Seave highlights the variant region in the UCSC genome browser. Seave can rapidly navigate to the site of each variant in an existing IGV (Robinson et al. 2011) session using the optional IGV column in the results table. Above the results table is a search box that allows immediate dynamic restriction of variant rows to any free text present anywhere in the results, e.g. entering “CDKN2A” or a disease name will only show variants where the gene or disease are mentioned anywhere in the variant’s annotations. Results are cached so that a URL of the results page can be shared with collaborators to let them see the same results page but without the ability to further query the database. Finally, all query options are saved so re-running analyses with modified parameters or inheritance models is a rapid process.

### Querying long variants

The Genome Block Store (GBS) within Seave stores long variants such as copy number variants, structural variants and losses of heterozygosity. GBS data can be queried either independently of short variants to explore the impact of just these events on the genome, or in conjunction with short variants to find sites of multiple hits. Similarly to short variants, GBS data is queried by families and samples, and several analysis types are available to explore the large variation in different ways. The GBS uses sample names to tie short variants to long variants and partially relies on BEDTools (Quinlan and Hall 2010) to perform interval querying logic.

#### Coordinate and gene list query

Querying by gene lists or genomic coordinates is the most common way to subset large variants to genomic regions with potential clinical impact. For example, a sample with a deletion spanning a gene of known disease association can be detected by querying the sample in the GBS for a gene list containing the gene. The tabular results from this query type contain rows for each gene or coordinate overlap associated with a separate large variant and sample.

#### Structural variant fusions query

The SV fusions analysis type interrogates translocation and inversion events that interrupt normal transcripts by fusing a gene to a distant genomic location. The breakpoints of these events are stored as linked blocks in the GBS and this analysis type finds all pairs of breakpoints where at least one is inside a gene of interest. The tabular results contain one fusion per row, called by a method for a sample, and include the gene(s) impacted along with the coordinates and annotations for each breakpoint.

#### Regions of homozygosity query

The GBS can store large regions of homozygosity identified by our in-house tool ROHmer (Puttick et al, manuscript in preparation). The ROHmer GBS analysis type identifies genomic regions of homozygosity that are shared by all affected individuals in a family but not shared by any unaffected individuals. These regions can be used to restrict the genomic search space to identify homozygous recessive variants that are the products of consanguinity.

#### Overlapping blocks query

Many groups run more than one CNV or SV caller, so the GBS allows the identification of all blocks that overlap between methods for the same sample. This is useful for determining regions of concordance between multiple variant callers. The GBS also allows finding overlapping blocks between samples for the same method which is useful for determining where multiple samples have shared variants called using the same method.

#### Short variants inside long variants

If a sample has both short variants and long variants in the GBS, they are automatically linked and returned in the short variant query results. This allows inferring second hit events such as a deletion combined with a damaging short variant or an inherited germline variant that has been duplicated to result in a gain of function. Seave does not currently include long variants during inheritance filtering so query results must be interpreted in light of Mendelian inheritance, e.g. a heterozygous short variant will become homozygous if there is an overlapping deletion and will no longer be returned for a heterozygous dominant inheritance filter.

### Somatic analyses

While Seave was initially created for germline rare disease analyses, it has been successfully used to interrogate somatic variants from multiple types of cancer sequencing using panels and whole genomes. Somatic variants called by Strelka (Saunders et al. 2012) or VarDict (Lai et al. 2016) have been used to create GEMINI (Paila et al. 2013) databases and imported into Seave for short variant analysis. Likewise, copy number variants from Sequenza (Favero et al. 2014), ClinSV (Minoche et al, in preparation) (genomes) or CNVkit (Talevich et al. 2016) (panels) and structural variants from Manta (Chen et al. 2016) can be imported into Seave’s GBS to enable CNV and fusions analysis.

We recommend that the germline and all tumour samples from the same patient are grouped into a single family, and that inheritance filters are skipped, to ignore the zygosity of any variant. The ability to group multiple samples together allows the investigation of changes in variants across multiple sites or through time within a single patient. We highlight variant allele frequency (VAF) instead of genotype when comparing across samples, using the overlapping CNVs and the purity estimate to infer clonality changes. Linking short and long variants allows rapidly inferring the mechanism by which a given somatic short variant has gained a foothold within the sample.

### Gene list management

Seave contains a custom gene list management system. Gene lists are uniquely named and have any number of genes associated with them. Upon addition to a gene list, all gene symbols are validated against a list of genes that can possibly be in Seave, based on the bioinformatics pipeline utilised. We recommend updating this list to the corresponding cache utilised by the version of VEP or snpEFF used to annotate VCFs input into Seave. We include the genes from the Ensembl 75 release, as this is the last to map to the b37 reference genome. Gene lists are available to all users within Seave for short and long variant query and can be modified at any time by administrators.

### Variant annotations and updates

Variant annotations place variants in the context of the functional genome; they are important ways of highlighting known or predicted important variants. Given the distributed nature of this information, annotations are often stored in separate databases with different structures at different locations. This makes determining the potential impact of a list of variants an arduous task, if undertaken manually. Seave aims to reduce this burden by storing local copies of a number of important annotations that are automatically joined to all short variant queries: OMIM (Amberger et al. 2015), ClinVar (Landrum et al. 2013), MITOMAP (Ruiz-Pesini et al. 2007), COSMIC (Forbes et al. 2010; 2015), COSMIC Cancer Gene Census (CGC) (Futreal et al. 2004), RVIS (Petrovski et al. 2013), MGRB (McNeil et al. 2017; 45 and Up Study Collaborators et al. 2008; MGRB 2017) and Orphanet (Orphanet 2017). We also include dbNSFP (Liu et al. 2011) which provides PROVEAN (Choi et al. 2012), FATHMM (Shihab et al. 2014), MetaSVM/MetaLR (Dong et al. 2015), GERP++ (Davydov et al. 2010) and Uniprot (The UniProt Consortium 2017) annotations. The annotations are stored in normalised MySQL databases to enable rapid variant annotation and can be updated as new versions of annotations are released. Using static freezes of annotations leads to out of date information and therefore missing important recent clinical relations between diseases and genetic variation. We provide code for updating and tracking annotations within Seave as they develop over time, which we run on a monthly basis.

### Data and user management

Seave caters to collaborative projects with a user and group management system. Variant data belongs to a group, and any number of users can belong to any number of groups. This makes segregating access to any given data set simple and shows every user all the data they have access to in one place. Administrator users are granted access to manipulate users, groups, variant data and gene lists in the administrator interface. All queries and changes to any aspect of Seave are logged for the purpose of keeping an audit trail.

### Deployment via AWS

To reduce the installation burden of Seave, we have created an Amazon Machine Image (AMI) that can be used to create a Seave server on Amazon Web Services (AWS) with ease. The AMI has been preconfigured to contain all dependencies that Seave requires and allows researchers to provision a server with desired specifications to match their needs. We recommend a t2.micro instance for a single group using Seave and a c4.large instance or above for an institution-scale Seave. AWS allows users to dynamically adjust resources if the workload changes or as the amount of data grows and provides simple administration options that allow encrypting genomic data and firewalling access to Seave to trusted IP ranges. By default, Seave is set up to run entirely on one server which includes an Apache webserver, MySQL database and space for a genomic data store. Alternatively, genomic data can be stored on an encrypted AWS Elastic Block Store (EBS) volume mounted on the Seave webserver and the MySQL database can be run on a dedicated AWS Relational Database Service (RDS) instance. A configuration file allows pointing the web-server to alternate locations easily, to the point where Seave can even be run locally.

### Testing and automation

Seave is developed using standard software engineering practices. Code development proceeds using continuous integration via git with automated testing and fixed deployments to various staging servers. Seave includes an end-to-end testing framework powered by Selenium which automatically performs a suite of variant queries on every code push and branch merge, to compare the results with what is expected and ensure no change to the code base has altered the results returned. We recommend those wishing to modify Seave to make use of these practices and contribute improvements back to the public repository via pull requests.

## Availability and licensing

To demo Seave with public data, navigate to: https://www.seave.bio. Seave is licensed exclusively for research or training use and the source code is available at: http://code.seave.bio. Extensive documentation is available at: http://documentation.seave.bio. For commercial and clinical diagnostic licensing, contact the corresponding author.

## Acknowledgements

We thank the Translational Genomics group (KCCG) and Genome.One for their comments and questions that lead to new features and bug fixes: Lisa Ewans, Eric Lee, Marie Wong, Kishore Kumar, Clare Puttick and Andr Minoche.

Seave would not be possible without the existence of GEMINI, a free academic tool created by Dr. Aaron Quinlan at the University of Utah. His work allowed us to rapidly prototype Seave and use GEMINI in any way we desire due to his generous MIT Licensing.

## Author contributions

VG and MJC conceived of and designed the software. TR provided clinical input. VG wrote the software. VG tested, debugged, designed individual features, and wrote documentation. MJC performed code review and developed the feature roadmap with VG. VG and MJC wrote the manuscript with input from all authors. MJC and MD supervised the project. All authors read and approved the final manuscript.

## Disclosure declaration

MED is the CEO of Genome.One Pty Ltd, which uses Seave as part of its diagnostic workflow. The remaining authors declare that they have no competing interests.

## Funding

VG, TR, MED and MJC were supported by a philanthropic grant from the Kinghorn Foundation. MJC was also supported by an Early Career Fellowship from Cancer Institute NSW (13/ECF/1-46) and grant ECP 1-96 from the NSW Department of Health.

## Additional Files

**Supplementary Figure 1 — Seave screenshot: database selection page**

This page shows the available databases for query and information about them, including the number of samples and variants. Upon logging in, this page displays databases from all groups the user has access to.

**Supplementary Figure 2 — Seave screenshot: family and analysis selection page**

After clicking a database to query, this page optionally allows selecting a family within it to query. If a family is selected, further options to select an analysis type (i.e. inheritance pattern) appear.

**Supplementary Figure 3 — Seave screenshot: filtration options/query page**

The query page allows the user to specify filtration options to restrict the number of variants returned. Restrictions can be by genomic location, impact on genes, prevalence in control populations and sequencing quality.

**Supplementary Figure 4 — Seave screenshot: results page**

Variants passing filtration criteria are displayed on the results page. A small number of default summary columns are immediately shown, highlighting the location and impact of the variant. The table is dynamic and additional annotation columns can be displayed by clicking any of the buttons under the table. The search box above the table searches across all columns to immediately restrict results to rows containing any terms entered (e.g. ‘missense variant chr10‘ to only see missense variants on chromosome 10). The GEMINI query used to generate the results can be displayed by clicking the “Show/hide GEMINI query” link under the table.

**Supplementary Figure 5 — Seave screenshot: results table expanded with additional columns**

The variants table on the results page can be expanded to show more annotation information. Any overflowing information can be read by hovering over the table cell and reading the tooltip that appears, as shown in this screenshot.

## References

1000 Genomes Project Consortium, Auton A, Brooks LD, Durbin RM, Garrison EP, Kang HM, Korbel JO, Marchini JL, McCarthy S, McVean GA, et al.. 2015. A global reference for human genetic variation. Nature Publishing Group 526: 68–74.

45 and Up Study Collaborators, Banks E, Redman S, Jorm L, Armstrong B, Bauman A, Beard J, Beral V, Byles J, Corbett S, et al.. 2008. Cohort profile: the 45 and up study. International journal of epidemiology 37: 941–947.

Abyzov A, Urban AE, Snyder M, and Gerstein M. 2011. CNVnator: an approach to discover, genotype, and characterize typical and atypical CNVs from family and population genome sequencing. Genome Research 21: 974–984.

Amberger JS, Bocchini CA, Schiettecatte F, Scott AF, and Hamosh A. 2015. OMIM.org: Online Mendelian Inheritance in Man (OMIM(R)), an online catalog of human genes and genetic disorders. Nucleic Acids Research 43: D789–D798.

Amendola LM, Jarvik GP, Leo MC, McLaughlin HM, Akkari Y, Amaral MD, Berg JS, Biswas S, Bowling KM, Conlin LK, et al.. 2016. Performance of ACMG-AMP Variant-Interpretation Guidelines among Nine Laboratories in the Clinical Sequencing Exploratory Research Consortium. American journal of human genetics 98: 1067–1076.

Balasubramaniam S, Riley LG, Bratkovic D, Ketteridge D, Manton N, Cowley MJ, Gayevskiy V, Roscioli T, Mohamed M, Gardeitchik T, et al.. 2017a. Unique presentation of cutis laxa with Leigh-like syndrome due to ECHS1 deficiency. Journal of inherited metabolic disease pp. 1–3.

Balasubramaniam S, Riley LG, Vasudevan A, Cowley MJ, Gayevskiy V, Sue CM, Edwards C, Edkins E, Junckerstorff R, Kiraly-Borri C, et al.. 2017b. EPG5-Related Vici Syndrome: A Primary Defect of Autophagic Regulation with an Emerging Phenotype Overlapping with Mitochondrial Disorders. JIMD reports.

Chen X, Schulz-Trieglaff O, Shaw R, Barnes B, Schlesinger F, Källberg M, Cox AJ, Kruglyak S, and Saunders CT. 2016. Manta: rapid detection of structural variants and indels for germline and cancer sequencing applications. Bioinformatics 32: 1220–1222.

Choi Y, Sims GE, Murphy S, Miller JR, and Chan AP. 2012. Predicting the functional effect of amino acid substitutions and indels. PLoS ONE 7: e46688.

Cingolani P, Platts A, Wang LL, Coon M, Nguyen T, Wang L, Land SJ, Lu X, and Ruden DM. 2012. A program for annotating and predicting the effects of single nucleotide polymorphisms, SnpEff: SNPs in the genome of Drosophila melanogaster strain w(1118); iso-2; iso-3. Fly 6: 80–92.

Davydov EV, Goode DL, Sirota M, Cooper GM, Sidow A, and Batzoglou S. 2010. Identifying a high fraction of the human genome to be under selective constraint using GERP++. PLoS Computational Biology 6: e1001025.

De Sousa SMC, McCabe MJ, Wu K, Roscioli T, Gayevskiy V, Brook K, Rawlings L, Scott HS, Thompson TJ, Earls P, et al.. 2017. Germline variants in familial pituitary tumour syndrome genes are common in young patients and families with additional endocrine tumours. European Journal of Endocrinology 176: 635–644.

Delaney SK, Hultner ML, Jacob HJ, Ledbetter DH, McCarthy JJ, Ball M, Beckman KB, Belmont JW, Bloss CS, Christman MF, et al.. 2016. Toward clinical genomics in everyday medicine: perspectives and recommendations. Expert review of molecular diagnostics 16: 521–532.

Dong C, Wei P, Jian X, Gibbs R, Boerwinkle E, Wang K, and Liu X. 2015. Comparison and integration of deleteriousness prediction methods for nonsynonymous SNVs in whole exome sequencing studies. Human molecular genetics 24: 2125–2137.

Eberle MA, Fritzilas E, Krusche P, Källberg M, Moore BL, Bekritsky MA, Iqbal Z, Chuang HY, Humphray SJ, Halpern AL, et al.. 2017. A reference data set of 5.4 million phased human variants validated by genetic inheritance from sequencing a three-generation 17-member pedigree. Genome Research 27: 157–164.

Favero F, Joshi T, Marquard AM, Birkbak NJ, Krzystanek M, Li Q, Szallasi Z, and Eklund AC. 2014. Sequenza: allele-specific copy number and mutation profiles from tumor sequencing data. Annals of Oncology 26: 64–70.

Forbes SA, Beare D, Gunasekaran P, Leung K, Bindal N, Boutselakis H, Ding M, Bamford S, Cole C, Ward S, et al.. 2015. COSMIC: exploring the world’s knowledge of somatic mutations in human cancer. Nucleic Acids Research 43: D805–11.

Forbes SA, Bindal N, Bamford S, Cole C, Kok CY, Beare D, Jia M, Shepherd R, Leung K, Menzies A, et al.. 2010. COSMIC: mining complete cancer genomes in the Catalogue of Somatic Mutations in Cancer. Nucleic Acids Research 39: D945–D950.

Futreal PA, Coin L, Marshall M, Down T, Hubbard T, Wooster R, Rahman N, and Stratton MR. 2004. A census of human cancer genes. Nature reviews. Cancer 4: 177–183.

Garrison E and Marth G. 2012. Haplotype-based variant detection from short-read sequencing. arXiv.org p. 1207.3907.

Heimer G, Kerätär JM, Riley LG, Balasubramaniam S, Eyal E, Pietikäinen LP, Hiltunen JK, Marek-Yagel D, Hamada J, Gregory A, et al.. 2016. MECR Mutations Cause Childhood-Onset Dystonia and Optic Atrophy, a Mitochondrial Fatty Acid Synthesis Disorder. The American Journal of Human Genetics 99: 1229–1244.

Kircher M, Witten DM, Jain P, O‘Roak BJ, Cooper GM, and Shendure J. 2014. A general framework for estimating the relative pathogenicity of human genetic variants. Nature Publishing Group 46: 310–315.

Kumar KR, Wali GM, Kamate M, Wali G, Minoche AE, Puttick C, Pinese M, Gayevskiy V, Dinger ME, Roscioli T, et al.. 2016. Defining the genetic basis of early onset hereditary spastic paraplegia using whole genome sequencing. Neurogenetics 17: 265–270.

Lai Z, Markovets A, Ahdesmaki M, Chapman B, Hofmann O, McEwen R, Johnson J, Dougherty B, Barrett JC, and Dry JR. 2016. VarDict: a novel and versatile variant caller for next-generation sequencing in cancer research. Nucleic Acids Research 44: e108.

Landrum MJ, Lee JM, Riley GR, Jang W, Rubinstein WS, Church DM, and Maglott DR. 2013. ClinVar: public archive of relationships among sequence variation and human phenotype. Nucleic Acids Research 42: D980–D985.

Layer RM, Chiang C, Quinlan AR, and Hall IM. 2014. LUMPY: a probabilistic framework for structural variant discovery. Genome Biology 15: R84.

Lek M, Karczewski KJ, Minikel EV, Samocha KE, Banks E, Fennell T, O‘Donnell-Luria AH, Ware JS, Hill AJ, Cummings BB, et al.. 2016. Analysis of protein-coding genetic variation in 60,706 humans. Nature Publishing Group 536: 285–291.

Liu X, Jian X, and Boerwinkle E. 2011. dbNSFP: a lightweight database of human nonsynonymous SNPs and their functional predictions. Human mutation 32: 894–899.

McKenna A, Hanna M, Banks E, Sivachenko A, Cibulskis K, Kernytsky A, Garimella K, Altshuler D, Gabriel S, Daly M, et al.. 2010. The Genome Analysis Toolkit: A MapReduce framework for analyzing next-generation DNA sequencing data. Genome Research 20: 1297–1303.

McLaren W, Gil L, Hunt SE, Riat HS, Ritchie GRS, Thormann A, Flicek P, and Cunningham F. 2016. The Ensembl Variant Effect Predictor. Genome Biology 17: 122.

McNeil JJ, Woods RL, Nelson MR, Murray AM, Reid CM, Kirpach B, Storey E, Shah RC, Wolfe RS, Tonkin AM, et al.. 2017. Baseline Characteristics of Participants in the ASPREE (ASPirin in Reducing Events in the Elderly) Study. The journals of gerontology. Series A, Biological sciences and medical sciences 72: 1586–1593.

Merlevede J, Droin N, Qin T, Meldi K, Yoshida K, Morabito M, Chautard E, Auboeuf D, Fenaux P, Braun T, et al.. 2016. Mutation allele burden remains unchanged in chronic myelomonocytic leukaemia responding to hypomethylating agents. Nature Communications 7: 1–13.

MGRB. 2017. Medical Genome Reference Bank. https://sgc.garvan.org.au/initiatives/mgrb.

Orphanet. 2017. The portal for rare diseases and orphan drugs. http://www.orpha.net.

Paila U, Chapman BA, Kirchner R, and Quinlan AR. 2013. GEMINI: Integrative Exploration of Genetic Variation and Genome Annotations. PLoS Computational Biology 9: e1003153–8.

Petrovski S, Wang Q, Heinzen EL, Allen AS, and Goldstein DB. 2013. Genic Intolerance to Functional Variation and the Interpretation of Personal Genomes. PLoS Genetics 9: e1003709–13.

Quinlan AR and Hall IM. 2010. BEDTools: a flexible suite of utilities for comparing genomic features. Bioinformatics 26: 841–842.

Riley LG, Cowley MJ, Gayevskiy V, Roscioli T, Thorburn DR, Prelog K, Bahlo M, Sue CM, Balasubramaniam S, and Christodoulou J. 2017. A SLC39A8 variant causes manganese deficiency, and glycosylation and mitochondrial disorders. Journal of inherited metabolic disease 40: 261–269.

Robinson JT, Thorvaldsdóttir H, Winckler W, Guttman M, Lander ES, Getz G, and Mesirov JP. 2011. Integrative genomics viewer. Nature Biotechnology 29: 24–26.

Ruiz-Pesini E, Lott MT, Procaccio V, Poole JC, Brandon MC, Mishmar D, Yi C, Kreuziger J, Baldi P, and Wallace DC. 2007. An enhanced MITOMAP with a global mtDNA mutational phylogeny. Nucleic Acids Research 35: D823–8.

Saunders CT, Wong WSW, Swamy S, Becq J, Murray LJ, and Cheetham RK. 2012. Strelka: accurate somatic small-variant calling from sequenced tumor-normal sample pairs. Bioinformatics 28: 1811–1817.

Shihab HA, Gough J, Mort M, Cooper DN, Day INM, and Gaunt TR. 2014. Ranking non-synonymous single nucleotide polymorphisms based on disease concepts. Human Genomics 8: 11.

Talevich E, Shain AH, Botton T, and Bastian BC. 2016. CNVkit: Genome-Wide Copy Number Detection and Visualization from Targeted DNA Sequencing. PLoS Computational Biology 12: e1004873.

Tan A, Abecasis GR, and Kang HM. 2015. Unified representation of genetic variants. Bioinformatics 31: 2202–2204.

Tennessen JA, Bigham AW, O‘Connor TD, Fu W, Kenny EE, Gravel S, McGee S, Do R, Liu X, Jun G, et al.. 2012. Evolution and functional impact of rare coding variation from deep sequencing of human exomes. Science 337: 64–69.

The UniProt Consortium. 2017. UniProt: the universal protein knowledgebase. Nucleic Acids Research 45: D158–D169.

